# Restoring Amyloid Clearance via Astrocytes: Z17 Is a Selective Inhibitor of CHI3L1 in Alzheimers Disease

**DOI:** 10.64898/2025.12.07.692801

**Authors:** Hossam Nada, Longfei Zhang, Baljit Kaur, Nelson García Vázquez, Moustafa Gabr

**Author notes:** To whom correspondence should be addressed: Moustafa T. Gabr.

## Abstract

Alzheimers disease (AD) is a progressive neurodegenerative disorder characterized by cognitive decline, accumulation of hallmark protein aggregates and substantial neuroinflammation. Chitinase-3-like protein 1 (CHI3L1) which is predominantly produced by activated astrocytes in the central nervous system (CNS). Overexpression of CHI3L1 has been implicated with AD progression and worsening of symptoms. Herein, we report the identification of **Z17** as a novel and selective CHI3L1 inhibitor which directly bind to CHI3L1 an equilibrium dissociation constant (*KD*) of 6.0 µM. In human iPSC-derived astrocytes, **Z17** acted as a dual-action regulator by reinstating astrocytic function and suppressing inflammation. Additionally, **Z17** rescued CHI3L1-induced impairment by dose-dependently restoring Aβ uptake and normalizing lysosomal proteolytic activity and pH. Furthermore, **Z17** effectively blocked CHI3L1-driven activation of the NF-κB pathway in human astrocytes, providing a mechanistic explanation for the functional rescue. The *in vitro* pharmacokinetic (PK) profiling of **Z17** demonstrated favorable drug-like properties for CNS development. These findings support the advancement of **Z17** as a selective CHI3L1 inhibitor capable of simultaneously mitigating neuroinflammation and restoring astrocytic clearance mechanisms, making it a highly promising therapeutic candidate for Alzheimers disease.

**Abstract Figure:** 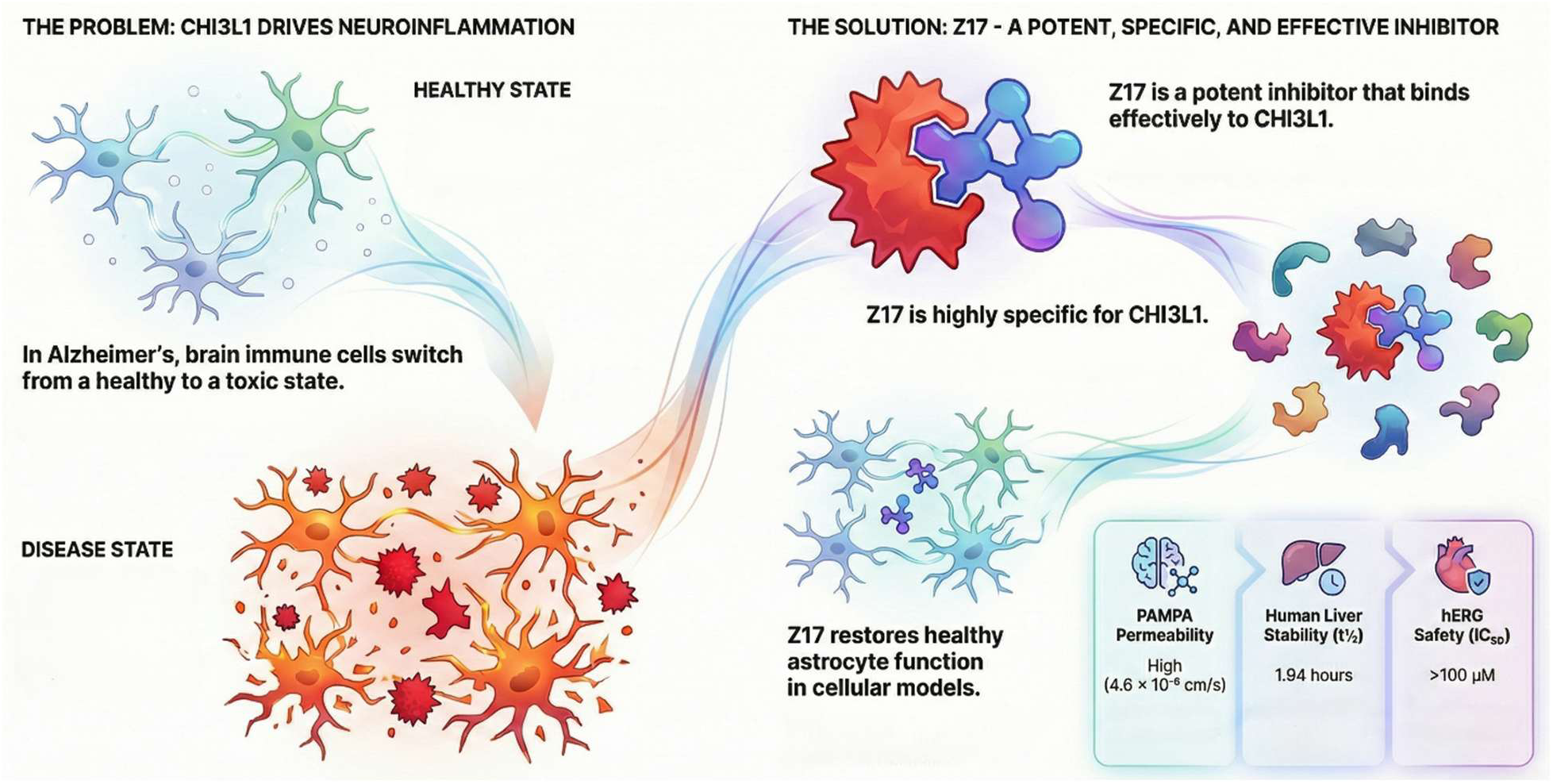

## 1. Introduction

Alzheimer’s disease (AD) is a neurodegenerative disorder affecting over 6.9 million^1^ individuals in the United States alone which is characterized by cognitive decline, memory impairment and ultimately patient mortality^2, 3^. Pathologically, AD is distinguished by the accumulation of extracellular amyloid-β (Aβ) plaques and intracellular neurofibrillary tangles composed of hyperphosphorylated tau protein, alongside substantial synaptic loss and neuronal death^4, 5^. While these hallmark protein aggregates have historically dominated AD research, emerging evidence increasingly implicates neuroinflammation as a central pathogenic mechanism rather than merely a secondary response to protein misfolding^5, 6^.

While neuroinflammation in the central nervous system (CNS) is essential for maintaining neuronal and brain homeostasis under physiological conditions, the transition of microglia and astrocytes into reactive, toxic states results in the dysregulation of neuroinflammation^7, 8^. Under normal circumstances, microglia mediate synaptic pruning and circuit refinement while astrocytes provide neuroprotection through clearance of toxic aggregates, release of trophic factors, and maintenance of the blood-brain barrier integrity^9, 10^. However, upon encountering pathogen-associated molecular patterns (PAMPs)^11^ or damage-associated molecular patterns (DAMPs)^12^ microglia and astrocytes release pro-inflammatory cytokines including IL-1β, IL-6, TNF-α, and chemokines such as CCL2 and CXCL1, alongside reactive oxygen species and nitric oxide (Figure 1). Persistent inflammatory activation results in continuous release of neurotoxic molecules, culminating in synaptic dysfunction, impaired neurogenesis, and progressive neurodegeneration^13, 14^.

**Figure 1.**
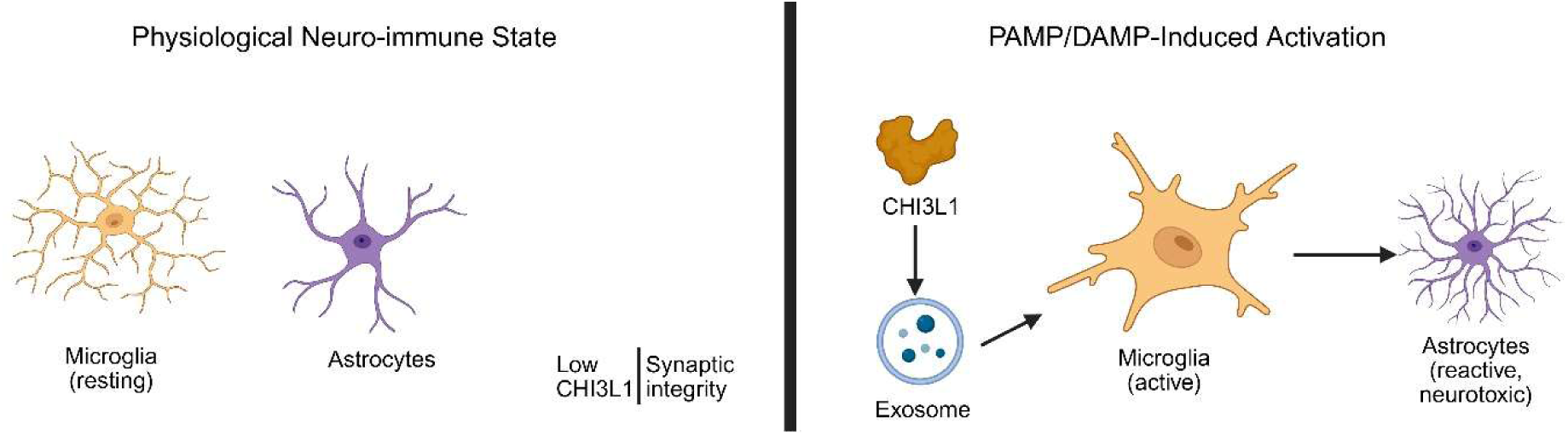
Resting versus activated neuroimmune states in the central nervous system. (A) Resting microglia and homeostatic astrocytes support synaptic stability and preserve neuronal health under physiological conditions, characterized by low CHI3L1 expression and minimal inflammatory activity. (B) Exposure to pathogen-associated molecular patterns (PAMPs) or damage-associated molecular patterns (DAMPs) activates microglia and drives astrocytes toward a reactive, neurotoxic A1 phenotype. This transition is accompanied by increased CHI3L1 production and exosome-mediated secretion, amplifying neuroinflammatory signaling.

Among the inflammatory mediators implicated in neurodegeneration, chitinase-3-like protein 1 (CHI3L1), also designated YKL-40 in humans based on its three N-terminal amino acids (tyrosine, lysine, leucine) and approximately 40 kDa molecular weight, has emerged as a key signaling molecule and promising biomarker^5, 15, 16^. Structurally, CHI3L1 contains an N-terminal chitin-binding domain homologous to the catalytic domain of chitinases and a C-terminal cysteine-rich domain involved in protein folding and stabilization through disulfide bond formation^17, 18^. Notably, despite its structural similarity to chitinases, CHI3L1 lacks enzymatic activity due to critical amino acid substitutions at the catalytic site^19, 20^. CHI3L1 undergoes extensive glycosylation, predominantly with complex-type N-glycans, which is critical for proper folding, stability, and secretion^5^. Unlike proteins utilizing classical secretion pathways, CHI3L1 lacks a conventional signal peptide and is instead released via non-classical mechanisms including exosome-mediated secretion and direct release from activated immune cells. In the extracellular space, CHI3L1 interacts with various matrix components including collagens, proteoglycans, and glycosaminoglycans, modulating cell-matrix adhesion, migration, and signaling pathways involved in tissue remodeling^5, 21, 22^.

In the CNS, CHI3L1 is predominantly produced by activated astrocytes and exhibits markedly elevated expression during neuroinflammatory conditions^22, 23^. Cerebrospinal fluid (CSF) and serum levels of CHI3L1 are consistently elevated in patients with multiple sclerosis, Parkinson’s disease, traumatic brain injury, and particularly AD, where levels correlate with disease severity, cognitive decline rate, and risk of disease development^24–26^. Remarkably, elevated CSF CHI3L1 can be detected in the earliest stages of AD, even preceding the onset of cognitive symptoms, and demonstrates a linear relationship with progression of cognitive impairment^27, 28^.

Despite extensive characterization of CHI3L1 functions in peripheral injury and repair responses, particularly in pulmonary inflammation where multiple receptors and downstream signaling pathways have been identified, the molecular mechanisms through which CHI3L1 mediates neuroinflammation and precipitates neurodegeneration in the brain remain poorly understood. Understanding how CHI3L1 functions as a signaling molecule in neuroinflammatory responses and how it contributes to the cascade of events leading to synaptic loss and cognitive decline in AD represents a critical knowledge gap. Consequently, the development of novel therapeutics capable of selectively modulating CHI3L1 expression or blocking its downstream signaling pathways represents a promising therapeutic avenue for mitigating neuroinflammation and improving clinical outcomes in AD patients.

Our previous efforts^29^ in identifying CHI3L1 inhibitors led to the discovery of two initial hits, **G28** and **E14**. However, both compounds presented limitations that prevented their advancement as viable lead candidates. **G28** demonstrated significant non-specificity, including a reported nanomolar affinity for human OX2R (IC₅₀ = 12 nM)^30^. In addition, SPR analysis of **G28** (Figure 4) revealed irregular binding profiles with aberrant association and dissociation kinetics, rendering it unsuitable for further development. **E14**, on the other hand, exhibited low potency, with a measured KD of 138 µM. Herein, we describe our optimization of E14, which led to the identification of **Z17**, a compound demonstrating a 20-fold improvement in potency. We further report the subsequent validation of **Z17** as a promising therapeutic candidate for Alzheimer’s disease.

## 2. Discussion

### 2.1. SAR exploration

To explore the structure–activity relationship (SAR) around **E4** and identify more potent CHI3L1 inhibitors, we initiated a focused analog-screening campaign. Commercially available compounds structurally related to **E4** were sourced from Enamine, prioritizing variations around the amide linkage, heterocyclic moieties, and substituent patterns on the aromatic core. This approach enabled rapid access to a diverse set of **E4**-like scaffolds without the need for de novo synthesis, allowing efficient assessment of key pharmacophoric features. A total of twenty-seven **E4** analogues (Figure 2) were procured and evaluated. These analogs captured modifications across the aliphatic side chain, ring size, electronic properties and steric bulk. Through this SAR exploration, several analogues, **Z5**, **Z9**, **Z10**, **Z17** and **Z20** displayed CHI3L1 binding activity.

**Figure 2.**
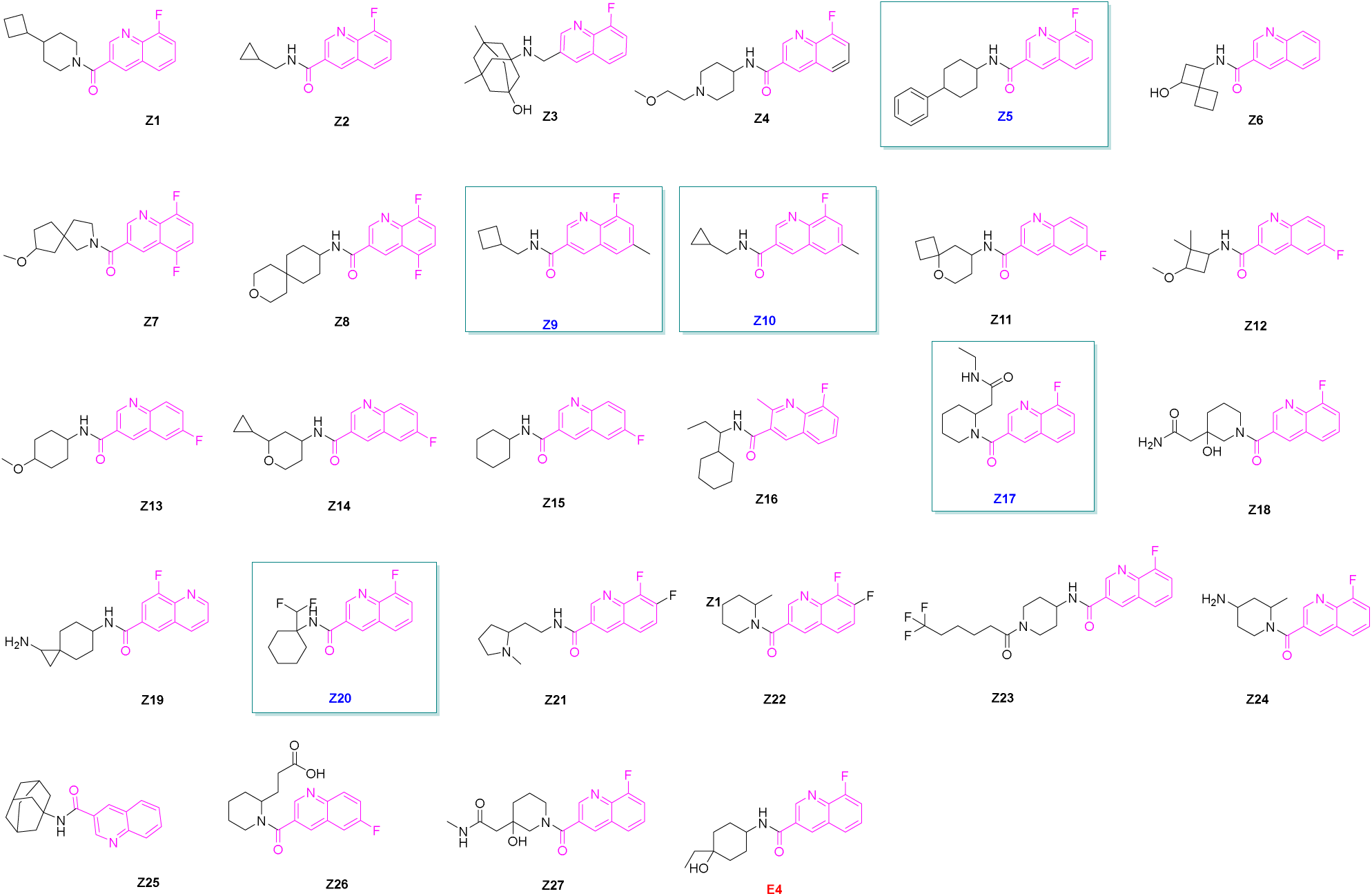
2D structures of E4 analogs explored for SAR-based optimization efforts.

### 2.2. Microscale Thermophoresis (MST) Single-Dose Screening

In the initial MST single-dose screen of 27 compounds, five compounds - **Z5**, **Z9**, **Z10**, **Z17** and **Z20** demonstrated detectable binding to CHI3L1 (Figure 3A). Additional control experiments were performed on these five initials to assess potential quenching and autofluorescence (Figure S1A-E and S1F). None of the tested compounds exhibited artifacts under these control conditions.

**Figure 3.**
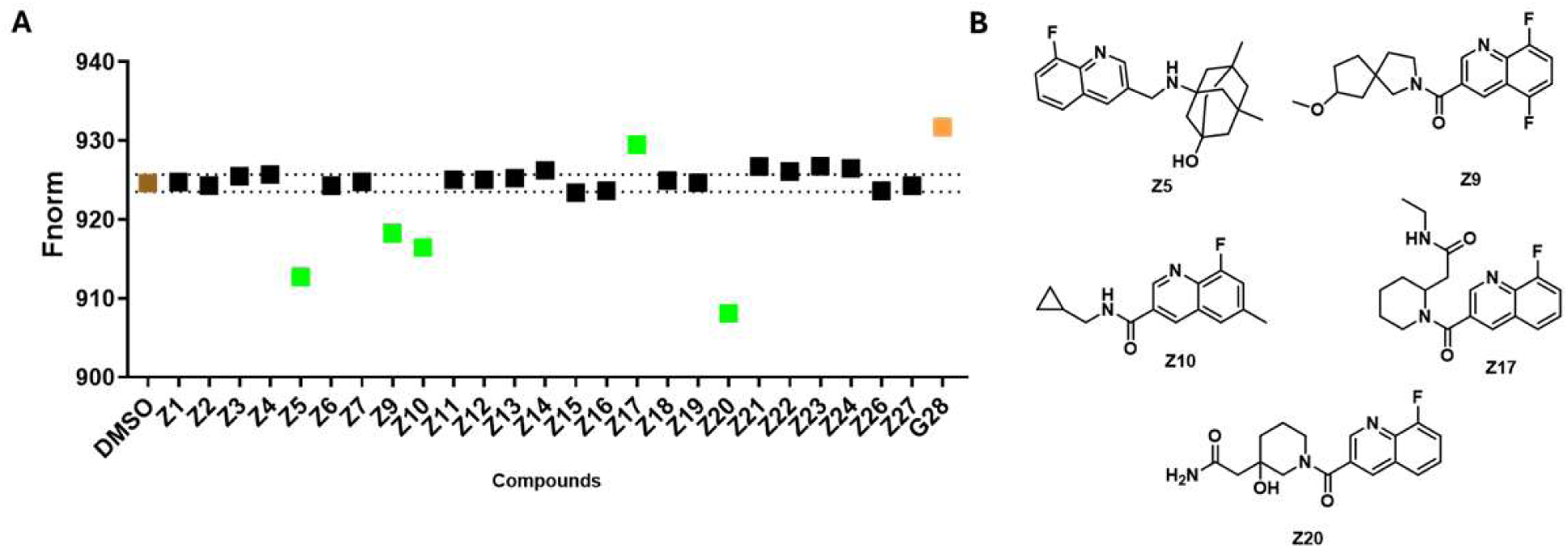
(A) Primary screening of compounds at 250 μM (with 2.5% DMSO). Green, brown, black, and orange indicate potential hits, negative control/reference (buffer with 2.5% DMSO), non-binders, and positive control (G28), respectively; (B) Chemical structures of the hit compounds.

**Figure 4.**
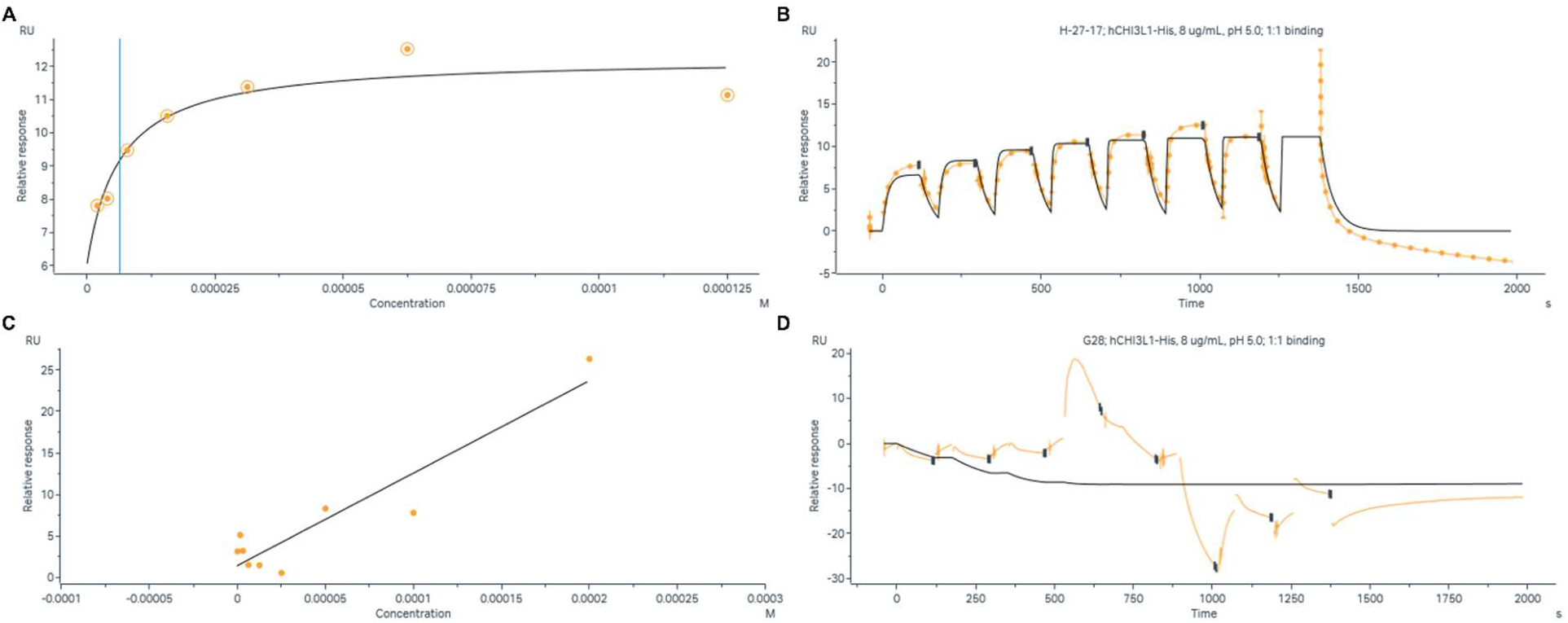
Comparative surface plasmon resonance (SPR) analysis of Z17 and G28 binding to hCHI3L1-His. (A) Steady-state affinity analysis showing the concentration-dependent binding of **Z17** to immobilized hCHI3L1-His protein. (B) Representative sensorgram showing real-time binding kinetics of **Z17** to hCHI3L1-His (8 µg/mL) at pH 5.0 under 1:1 binding stoichiometry. (C) Steady-state analysis of **G28** binding to hCHI3L1-His showing poor correlation with the 1:1 binding model. (D) Representative sensorgram for **G28** binding to hCHI3L1-His showing irregular binding profiles with aberrant kinetics. All experiments were performed at 25°C using PBS-P+ supplemented with 2.5% DMSO at pH 5.0.

### 2.3. SPR

The five hits from the single-dose screening were subjected to Surface plasmon resonance (SPR) assay to determine their dose response. The remaining four initial hits did not show a dose-dependent response. Surface plasmon resonance (SPR) analysis was employed to quantify the binding affinity of **Z17** to hCHI3L1-His at pH 5.0 under 1:1 binding stoiometry conditions. The equilibrium dissociation constant (K_D_) was determined to be 6.0 µM, indicating potent binding affinity between **Z17** and CHI3L1 (Figure 4). The binding kinetics sensorgram revealed characteristic association and dissociation phases across multiple analyte concentrations (8 µg/mL hCHI3L1-His), with rapid association kinetics observed during the injection phase followed by measurable dissociation upon buffer flow. The steady-state affinity analysis demonstrated a hyperbolic binding isotherm with saturation observed at higher concentrations, yielding a maximum response (R_max_) of approximately 12 RU. The micromolar affinity observed for **Z17** is consistent with small molecule inhibitors targeting protein-protein interaction interfaces and suggests that Z17 engages hCHI3L1 through a reversible, concentration-dependent mechanism.

SPR analysis of the reference compound **G28** revealed substantially weaker and inconsistent binding characteristics (Figure 4C-D). The steady-state analysis for **G28** (Figure 4C) showed poor correlation with the 1:1 binding model, with significant scatter in the data points and a response that increased approximately linearly rather than exhibiting saturable binding kinetics. The corresponding sensorgram (Figure 4D) displayed irregular binding profiles with aberrant association and dissociation kinetics, including negative response values and poor fit to the binding model (black line), suggesting non-specific interactions or compound aggregation artifacts.

### 2.4. Specificity assay

To evaluate the specificity of Z17, binding interactions with different proteins were examined using microscale thermophoresis (MST). Seven proteins were tested: triggering receptor expressed on myeloid cells 2 (TREM-2), roundabout homolog 1 (ROBO-1), cluster of differentiation 28 (CD28), inducible T-cell co-stimulator (ICOS), lymphocyte activation gene 3 (LAG-3), cluster of differentiation 80 (CD80), and galectin-3 (GAL-3).

Different buffer conditions were used depending on the protein as reported in literature. For TREM-2, ROBO-1, CD28, ICOS, LAG-3, and CD80, PBST buffer containing 0.05% Tween-20 was used. For ICOS, an alternative assay buffer consisting of 10 mM HEPES, 150 mM NaCl, and 0.005% Tween-20 was used. For GAL-3, the same assay conditions as those used for chitinase-3-like protein 1 (CHI3L1) were followed. In all cases, the protein concentration was maintained at 40 nM, while the compound concentration was kept at 100 μM with a final DMSO concentration of 2.5%. Same procedure as described for the initial screening of the compound library with CHI3L1 was used. Interestingly, **G28** displayed binding affinity toward GAL-3 (Figure 5G). However, **Z17** showed no detectable binding to any of the tested proteins, confirming the high selectivity of this compound (Figure 5A-3G). When combined with the fact that **G28** was previously reported to bind human OX_2_R with nanomolar affinity (IC₅₀ = 12 nM)^30^. The marked improvement in specific CHI3L1 binding observed with **Z17** compared to **G28**, coupled with Z17’s lack of OX2R activity, demonstrates successful structure-activity optimization toward a selective CHI3L1 inhibitor with well-defined target engagement and superior therapeutic potential.

**Figure 5.**
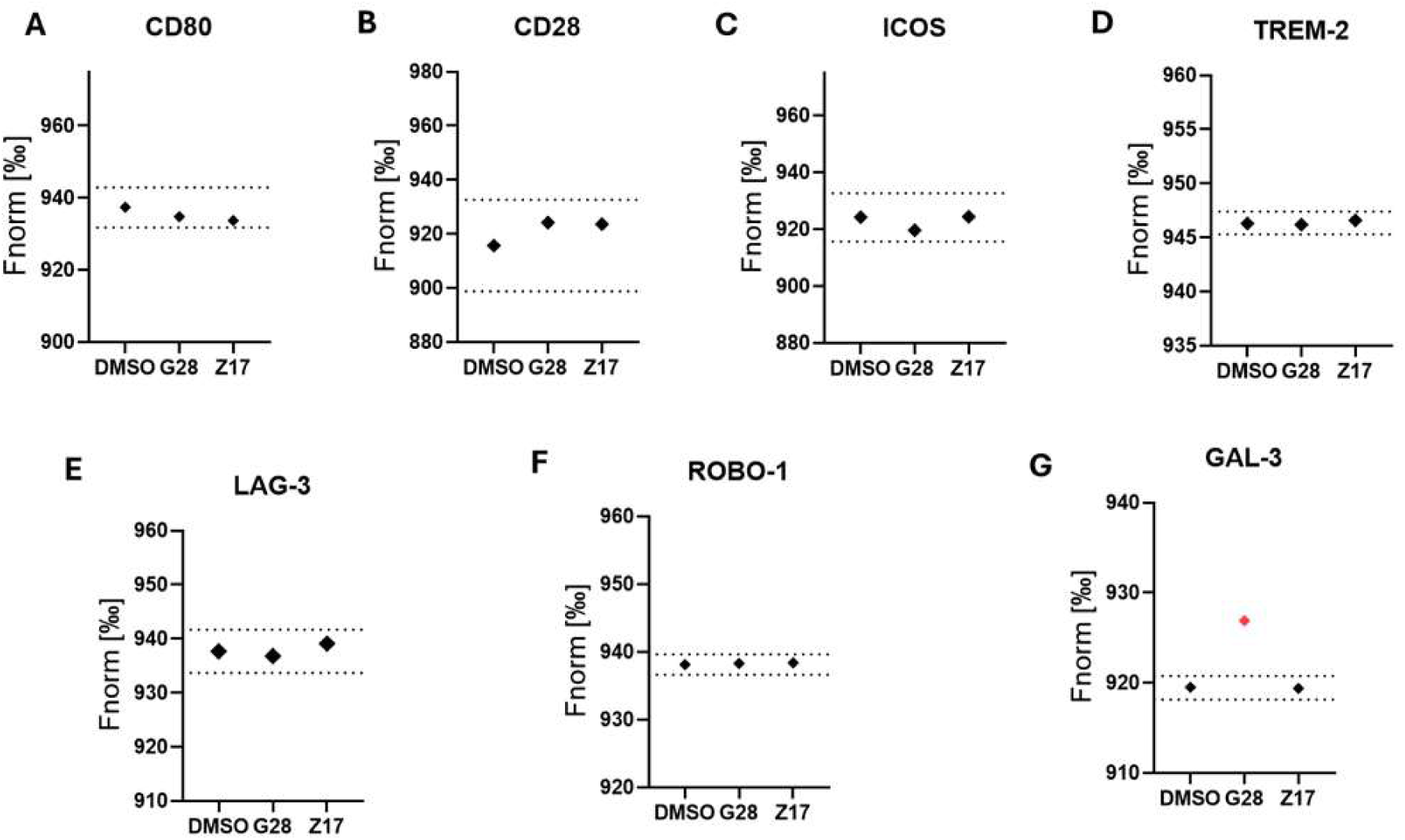
Specificity assessment of Z17 and G28 with different proteins.

### 2.5. In vitro PK Profiling of Z17

Advancing CNS-active small molecules for AD requires compounds with physicochemical and PK characteristics compatible with oral dosing, brain penetration, and chronic safety. To determine whether **Z17** meets these criteria, we carried out a comprehensive panel of in vitro ADME assays (Table 1).

**Table 1.** In vitro PK profile of Z17.

| PK property | Value |
| --- | --- |
| LogD7.4<sup>a</sup> | 2.39 ± 0.22 |
| Kinetic Solubility 1% DMSO/PBS [μM]<sup>b</sup> | 85 ± 10 |
| PAMPA permeability [cm/s] | 4.6 × 10<sup>-6</sup> |
| PPB human [%]<sup>c</sup> | 94.2 ± 0.6 |
| t<sub>1/2</sub> [h] mouse liver microsomes<sup>c</sup> / CL<sub>int</sub><sup>d</sup> | 1.52 ± 0.18 / 28 ± 4.1 |
| t<sub>1/2</sub> [h] human liver microsomes<sup>c</sup> / CL<sub>int</sub><sup>d</sup> | 1.94 ± 0.21 / 15 ± 2.6 |
| t<sub>1/2</sub> [h] mouse plasma | 3.1 ± 0.2 |
| $t_{1/2}$ [h] human plasma | 3.4 ± 0.3 |
| $t_{1/2}$ [h] PBS | > 6 |
| IC<sub>50</sub> against hERG<br>[μM] | >100 |
| IC<sub>50</sub> against WI-38<br>[μM] | >100 |
| IC<sub>50</sub> against HS 27<br>[μM] | >100 |
| IC<sub>50</sub> against NHA<br>[μM] | >100 |

<sup>a</sup>LogD (pH = 7.4) the distribution coefficient at physiological pH 7.4. <sup>b</sup>Kinetic solubility measured in 1% DMSO/PBS (pH 7.4) by UV-vis spectrophotometry (n=3). <sup>c</sup>Values are mean ± SD (n=3). <sup>d</sup>Intrinsic clearance expressed in μL/(min × mg) protein. Values are mean ± SD (n=3).

**Z17** showed a LogD_7.4_ of 2.39 ± 0.22, indicating a balanced hydrophobic-hydrophilic profile consistent with efficient intestinal absorption and favorable BBB permeability. Molecules within this lipophilicity range typically demonstrate robust CNS penetration while minimizing transporter-mediated efflux. **Z17** also exhibited a kinetic solubility of 85 ± 10 µM in PBS (1% DMSO), supporting the feasibility of achieving therapeutic free-drug concentrations without specialized formulation efforts. Passive permeability, evaluated using PAMPA, was high (4.6 × 10^-6^ cm s⁻¹), placing **Z17** well within the permeability range associated with passive transcellular BBB transport. These data are consistent with the structure’s modest polarity and tertiary amine functionality, both of which are hallmarks of CNS-penetrant chemotypes.

Microsomal stability studies indicated moderate intrinsic clearance (Table 1). **Z17** displayed half-lives of 1.52 ± 0.18 h in mouse microsomes and 1.94 ± 0.21 h in human microsomes, corresponding to CL_int_ values of 28 ± 4.1 µL min⁻¹ mg⁻¹ and 15 ± 2.6 µL min⁻¹ mg⁻¹, respectively. The improved stability of **Z17** in human microsomes is consistent with a reduced metabolic turnover in human liver enzymes and suggests favorable human PK translation.

**Z17** also demonstrated excellent plasma stability, with half-lives of 3.1 ± 0.2 h (mouse) and 3.4 ± 0.3 h (human), and remained intact for over 6 h in PBS, confirming that enzymatic pathways rather than chemical decomposition dominate its clearance (Table 1). Plasma protein binding was 94.2 ± 0.6%, typical for CNS drug candidates and supportive of sustained systemic exposure with an adequate unbound fraction for CNS target engagement. No measurable inhibition of the hERG channel was observed up to 100 µM, reducing potential liabilities related to cardiac electrophysiology. Similarly, **Z17** showed IC_50_ > 100 µM in WI-38, HS-27, and normal human astrocyte viability assays, indicating a wide in vitro safety margin (Table 1).

Together, these results (Figure 6, Table 1) indicate that **Z17** possesses a PK and safety profile well-suited for CNS drug development, with properties predictive of robust oral bioavailability, efficient BBB permeation, and a half-life compatible with once-or twice-daily dosing. These findings support advancement of **Z17** into in vivo PK and efficacy studies to confirm brain exposure and evaluate its therapeutic potential.

**Figure 6.**
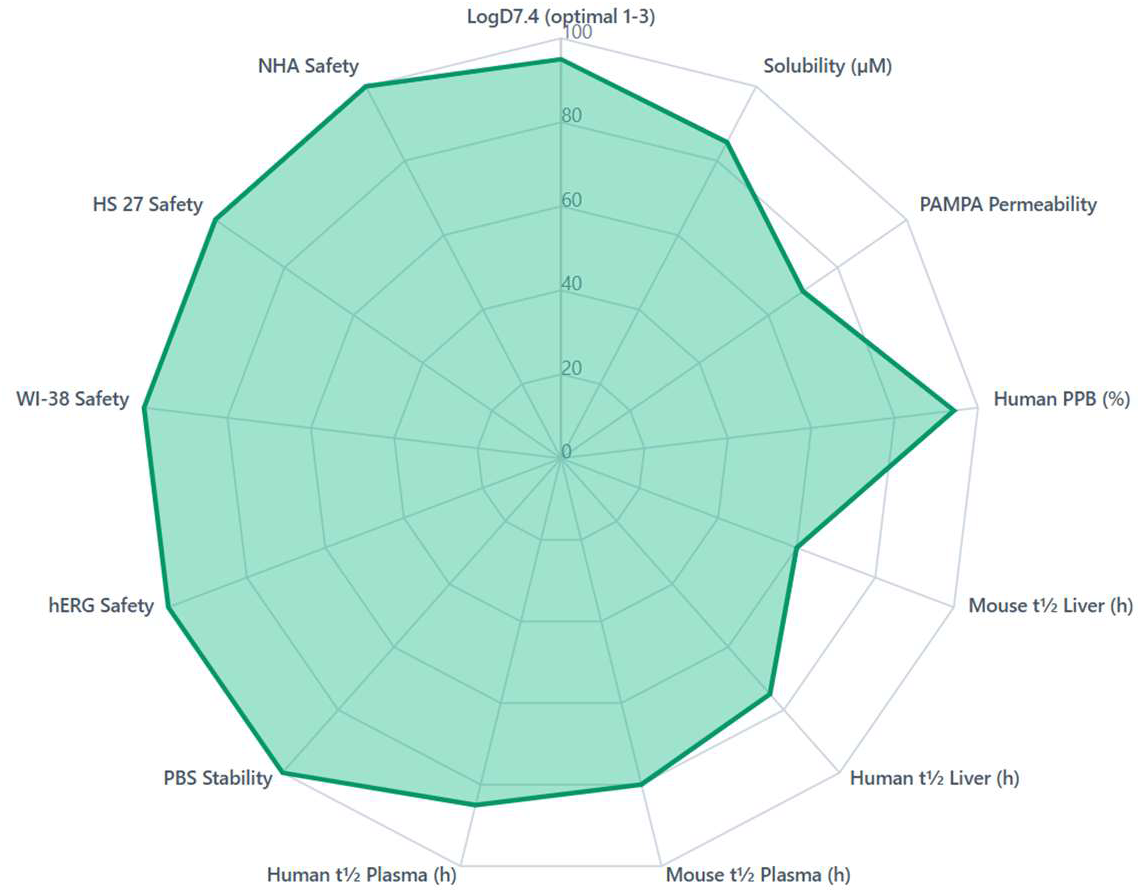
Spider diagram representation of the in vitro pharmacokinetic profile of compound Z17. All parameters are normalized to a desirability score ranging from 0 to 100, where higher values represent more favorable drug-like properties.

### 2.7. In vitro PK Profiling of Z17

#### 2.7.1. **Z17** Restores Aβ Uptake and Lysosomal Function in CHI3L1-Stimulated Astrocytes

To determine whether **Z17** can counteract CHI3L1-mediated astrocyte dysfunction, we evaluated its effects on endocytic efficiency, lysosomal degradation, and pH homeostasis in human iPSC-derived astrocytes exposed to recombinant CHI3L1. Consistent with prior reports, CHI3L1 (300 ng/mL) markedly impaired uptake of pHrodo-Aβ_1-42_, reducing internalization to 0.63-fold of vehicle-treated levels. Pre-treatment with **Z17** improved uptake in a clear dose-responsive manner, where 50 µM of **Z17** fully rescued the phenotype (Figure 7A). CHI3L1 also suppressed lysosomal proteolytic activity, as measured by DQ-BSA degradation, decreasing fluorescent signal to 0.61-fold of vehicle. **Z17** reversed the deficit, elevating degradative capacity to 0.72 ± 0.03-fold at 10 µM, 0.83 ± 0.04-fold at 25 µM, and 0.95 ± 0.11-fold at 50 µM (Figure 7B). In parallel, the CHI3L1-induced increase in LysoSensor ratio (from 1.00 to 1.43 ± 0.09) was normalized by **Z17**. The compound reduced the elevated ratio to 1.35 ± 0.03 at 10 µM, 1.23 ± 0.04 at 25 µM, and 1.09 ± 0.05 at 50 µM, indicating restored lysosomal acidity (Figure 7C).

**Figure 7.**
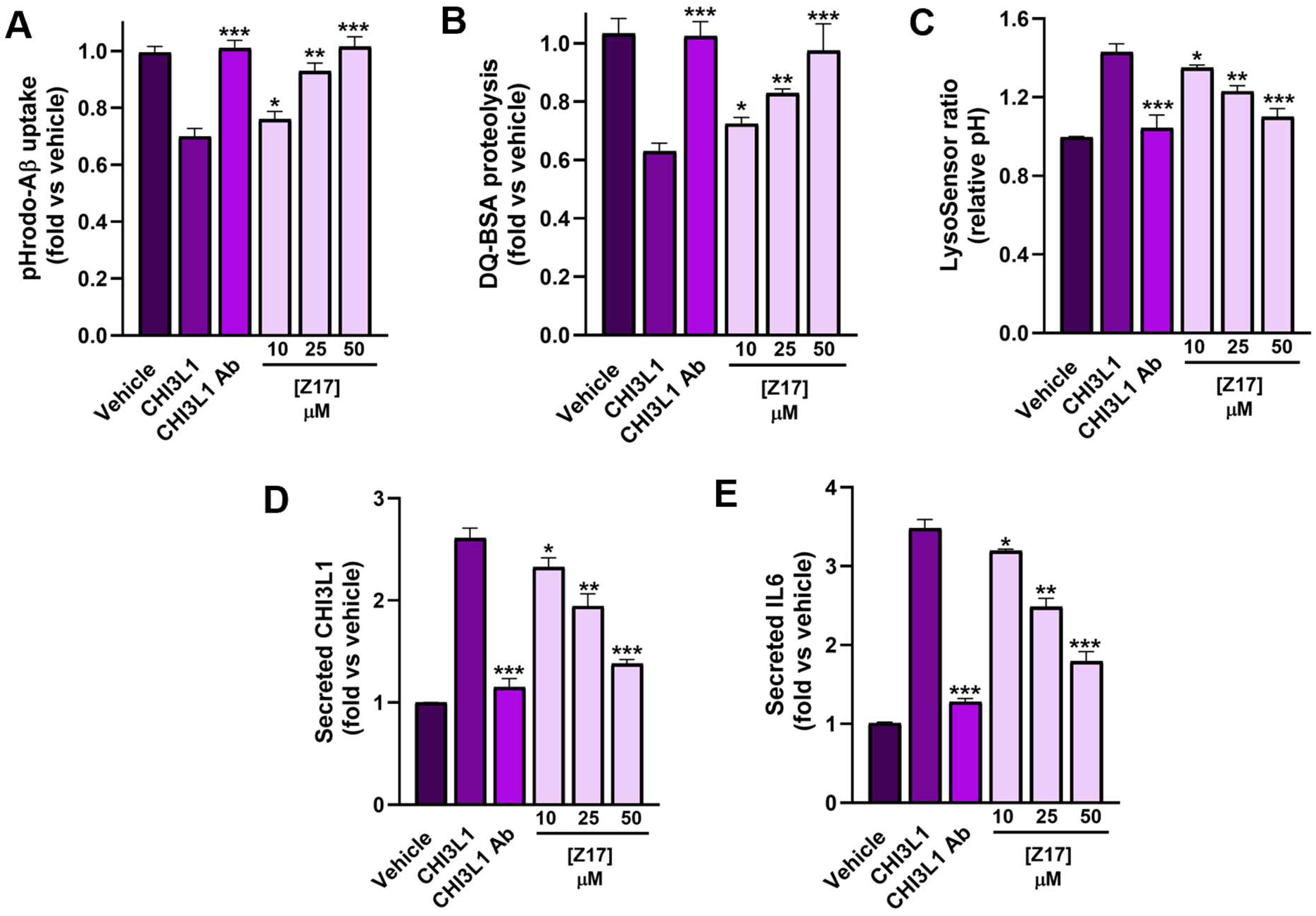
Z17 restores Aβ uptake, lysosomal function, and cytokine balance in CHI3L1-treated astrocytes. Recombinant CHI3L1 (300 ng/mL) reduced pHrodo-Aβ_1-42_ uptake in human iPSC-derived astrocytes to 68 ± 6% of vehicle, whereas **Z17** restored uptake in a dose-dependent manner at 10, 25, and 50 µM **(A)**. CHI3L1 impaired lysosomal proteolysis (DQ-BSA) and increased lysosomal pH, and both defects were corrected by **Z17** across the same concentration range **(B-C)**. **Z17** also reduced CHI3L1-stimulated secretion of CHI3L1 and IL-6 without affecting cell viability **(D-E)**. The neutralizing anti-CHI3L1 antibody (10 µg mL^-1^) was used as a positive control in all assays. Data represent mean ± SD from n = 3 experiments. Statistical analysis was performed by one-way ANOVA with Dunnett’s post hoc test versus CHI3L1: *p* < 0.05 (\**), p < 0.01 (*\*\**), p < 0.001 (*\*\*\*).

**Z17** also dampened the cytokine response triggered by CHI3L1, lowering the secretion of CHI3L1 itself as well as IL-6, without compromising astrocyte viability (Figures 7D,E). Together, these data show that **Z17** counteracts CHI3L1-mediated astrocyte impairment by restoring lysosomal proteolytic function and normalizing inflammatory signaling. By directly engaging CHI3L1, **Z17** disrupts the aberrant CHI3L1-CRTH2/RAGE pathway, blocking downstream IKKβ–S6K1-S6 activation and maintaining proper vesicular pH. This rescue phenotype mirrors that of CHI3L1-deficient astrocytes, which display elevated Aβ uptake and reduced IL-6 release. Overall, the findings establish **Z17** as a dual-action regulator that simultaneously suppresses CHI3L1-driven neuroinflammation and reinstates astrocytic Aβ clearance, two linked mechanisms implicated in AD progression.

#### 2.7.2. **Z17** Suppresses CHI3L1-Induced NF-κB Signaling in Human Astrocytes

To establish whether **Z17** directly interferes with the inflammatory signaling cascade initiated by CHI3L1, we assessed NF-κB transcriptional activity using a dual-luciferase reporter system in human iPSC-derived astrocytes. Consistent with its known role as a pro-inflammatory effector, recombinant CHI3L1 (300 ng/mL) markedly increased NF-κB-driven firefly luciferase activity, resulting in a 3.85 ± 0.2-fold induction relative to vehicle-treated controls. Preincubation with **Z17** produced a clear, concentration-dependent reduction in CHI3L1-induced NF-κB activation (Figure 8A). Exposure to 10 µM **Z17** decreased the reporter signal to 3.38 ± 0.1-fold, whereas 25 µM further reduced activation to 2.51 ± 0.4-fold. At 50 µM, **Z17** lowered NF-κB activity to 1.95 ± 0.1-fold, approaching the suppression achieved with a neutralizing anti-CHI3L1 antibody (10 µg/mL), which reduced signaling to 1.48 ± 0.2-fold.

**Figure 8.**
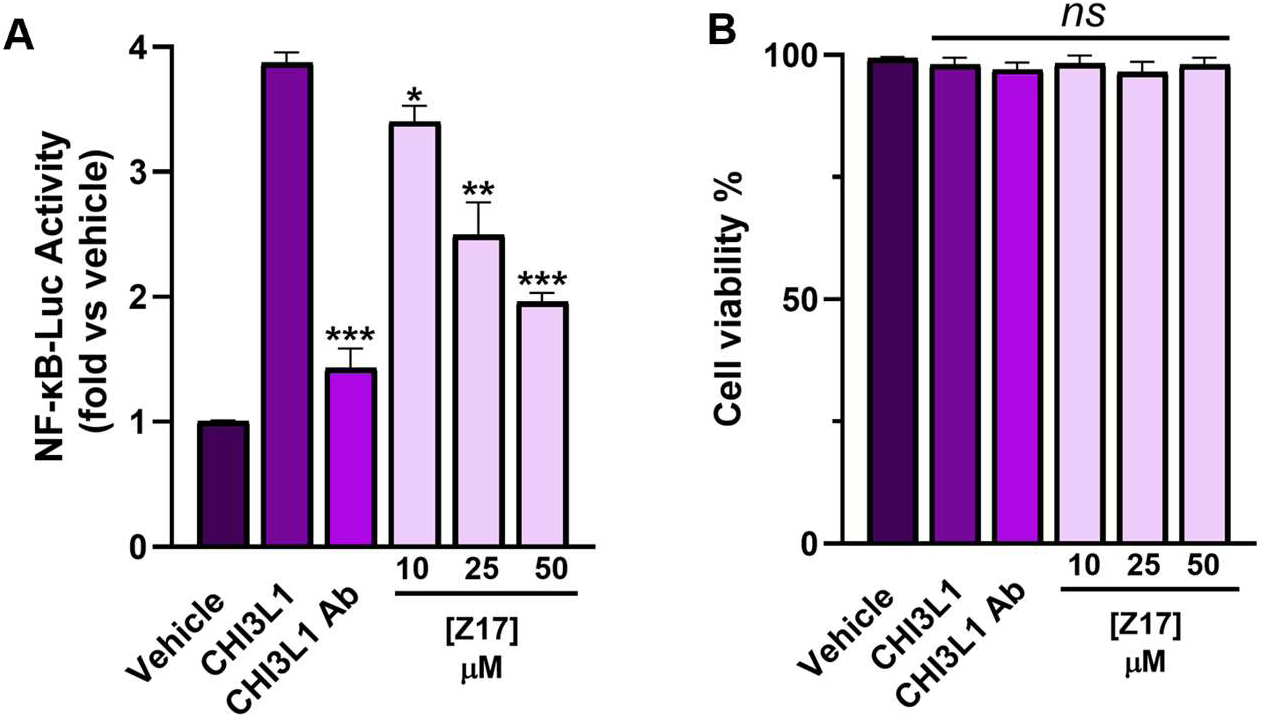
Z17 inhibits CHI3L1-induced NF-κB activation in human astrocytes. CHI3L1 (300 ng/mL) increased NF-κB-luciferase activity to 3.85 ± 0.2-fold over vehicle. **Z17** reduced this activation in a concentration-dependent manner at 10, 25, and 50 µM, with the highest dose approaching the effect of a neutralizing anti-CHI3L1 antibody (10 µg mL^-1^) **(A)**. Renilla normalization and viability assays confirmed that signal reduction was not due to cytotoxicity **(B)**. Data represent mean ± SD from n = 3 experiments. One-way ANOVA with Dunnett’s post hoc test versus CHI3L1: *ns*, not significant; *p* < 0.05 (\**), p < 0.01 (*\*\**), p < 0.001 (*\*\*\*).

Importantly, **Z17** did not alter Renilla luciferase activity or compromise cell viability across the concentration range tested, indicating that the reduction in NF-κB signal was not attributable to nonspecific cytotoxicity or assay interference (Figure 8B). These findings demonstrate that Z17 effectively blocks CHI3L1-driven activation of the NF-κB pathway in human astrocytes. Given the established link between CHI3L1 signaling, IKKβ phosphorylation, and downstream S6K1-mediated transcriptional activation, the inhibitory profile of **Z17** suggests a direct interruption of the CHI3L1-CRTH2/RAGE-IKKβ axis. Together with its restoration of lysosomal function and attenuation of cytokine secretion, the suppression of NF-κB activity provides a mechanistic explanation for the broad functional rescue exerted by **Z17** in CHI3L1-stimulated astrocytes.

## 3. Experimental

### 3.1. MST Screening

Protein–ligand interactions were assessed using MST following our previously published protocol. CHI3L1-His protein was fluorescently labeled with RED-tris-NTA dye (NanoTemper Technologies) according to the manufacturer’s instructions. Labeled protein was incubated with test compounds under established assay conditions and analyzed using the Dianthus NT.23 Pico system. To control for potential artifacts from autofluorescence and quenching, all the hits were subjected to additional pretests. For autofluorescence, fluorescence of labeled protein with 2.5% DMSO was compared to buffer containing 250 μM compound in 2.5% DMSO (Figure S1A-E). For quenching assessment, fluorescence from 20 nM dye in buffer with 2.5% DMSO was compared to that of 20 nM dye incubated with each compound in 2.5% DMSO (Figure S1F). Compounds outside the mean ± 3 SD of the reference were excluded; none of the tested compounds exhibited autofluorescence or quenching.

### 3.2. SPR

SPR dose-response analysis was performed on the five hit compounds identified from single-dose screening using a Biacore™ 8K system (Cytiva, Marlborough, MA, USA) to validate binding and determine equilibrium dissociation constants. hCHI3L1-His protein (8.0 μg/mL in acetate buffer, pH 5.0) was immobilized on a Series S Sensor Chip CM5 (29104988, Cytiva) via amine coupling using a commercial amine coupling kit (BR100050, Cytiva) at a flow rate of 10 μL/min for 420 s, achieving an immobilization level of approximately 3000 RU, followed by blocking with ethanolamine. For dose-response evaluation, serial dilutions of each compound were prepared in PBS-P+ (28995084, Cytiva) supplemented with 2.5% DMSO to achieve concentrations ranging from 0.78 to 100 μM. Test compounds were injected over the sensor chip using a multi-cycle kinetics protocol at a flow rate of 30 μL/min for 120 s (association phase), followed by buffer flow for 120 s (dissociation phase). Regeneration between cycles was performed using regeneration buffer (10 mM sodium acetate, 250 mM NaCl, pH 4.0) at 30 μL/min for 30 s. A reference flow cell blocked solely with ethanolamine on the same sensor chip was used for double referencing. Sensorgram data were analyzed using Biacore™ Insight Evaluation Software (Cytiva) with a 1:1 binding model to determine kinetic parameters (ka, kd) and equilibrium dissociation constants (KD).

### 3.3. CHI3L1 Stimulation and Compound Treatment

Recombinant CHI3L1 protein was used to activate signaling pathways in U-87 MG cells. To assess the inhibitory potential of compound Z17, CHI3L1 was pre-incubated with Z17 for 1 h at room temperature in serum-free RPMI 1640 to allow potential ligand–compound interaction before cell exposure. Following serum starvation, cells were mechanically detached to achieve uniform suspensions, and 4 × 10⁵ cells were distributed into 1.5 mL microcentrifuge tubes for each treatment condition.

Cells were treated with one of three conditions: (i) DMSO vehicle control, (ii) CHI3L1 + DMSO, or (iii) CHI3L1 + Z17. Treatments were carried out in serum-free RPMI 1640 for 15 min at 37 °C to capture early phosphorylation events downstream of receptor engagement. Cells were then pelleted by centrifugation, supernatant was removed, and pellets were lysed directly in RIPA buffer supplemented with protease and phosphatase inhibitors.

### 3.4. Western Blot Analysis

Total protein concentration was determined by bicinchoninic acid (BCA) assay. Equal amounts of lysate were resolved by SDS–PAGE and transferred to PVDF membranes. Membranes were blocked in 5% non-fat dry milk in TBS-T and incubated overnight at 4 °C with primary antibodies against phospho-ERK1/2 (Thr202/Tyr204), COX-2, and Vinculin (loading control). After washing, membranes were incubated with HRP-conjugated secondary antibodies and developed using enhanced chemiluminescence (ECL).

### 3.5. Evaluation of PK and physicochemical properties

The These experiments were conducted following our previously reported methods.^31^ The evaluations included LogD7.4 determination, microsomal stability, kinetic solubility, and cytotoxicity profiling across multiple cell lines. Solubility was assessed using UV–visible spectrophotometry, while cell viability was measured using the PrestoBlue assay.

### 3.6. Cell culture and reagents

Human iPSC-derived astrocytes were maintained according to the manufacturer’s protocol in astrocyte medium (DMEM/F12 supplemented with 10% FBS) at 37 °C in a humidified atmosphere of 5% CO₂. Recombinant human CHI3L1 (YKL-40) was purchased from R&D Systems. For comparison and pathway validation, a neutralizing anti-CHI3L1 antibody (R&D Systems) was used as a positive control. All assays were performed in 96-well format unless otherwise indicated.

### 3.7. Astrocyte Aβ Uptake and Lysosomal Function Assay

Human iPSC-astrocytes were seeded at 2 × 10^4^ cells per well (96-well plates) and cultured for 48 h. Cells were pre-treated with **Z17** (10-50 µM, 30 min) prior to stimulation with CHI3L1 (300 ng mL⁻¹, 24 h). For Aβ uptake, cells were incubated with pHrodo™-Red–labeled Aβ_1-42_ (1 µM) for 2 h, dissociated using Accutase, fixed in 2% paraformaldehyde, and analyzed on a BD Fortessa flow cytometer (Ex/Em 560/585 nm). The mean fluorescence intensity (MFI) from ≥10,000 live events per well was normalized to vehicle controls.

Lysosomal degradation was assessed using DQ-BSA Green (10 µg mL⁻¹, 2 h), and lysosomal pH was measured using LysoSensor™ DND-160 (Ex/Em 340/440–535 nm ratio). Supernatants were collected for CHI3L1 and IL-6 quantification by ELISA (R&D Systems). Cell viability was assessed using CellTiter-Glo® (Promega). All measurements were performed in triplicate from at least three independent experiments.

### 3.8. NF-κB Reporter Assay

To evaluate the effect of **Z17** on CHI3L1-driven inflammatory signaling, human iPSC-astrocytes were transduced with an NF-κB-firefly luciferase reporter and a Renilla luciferase internal control. Cells were maintained for 72 h post-transduction to achieve stable expression before treatment. Cells were pre-incubated with **Z17** (10-50 µM, 30 min) followed by stimulation with CHI3L1 (300 ng mL⁻¹, 6 h). Firefly and Renilla luciferase activities were measured using the Dual-Glo® Luciferase Assay System (Promega). Results were expressed as normalized Firefly/Renilla ratios and presented as fold-change relative to untreated vehicle controls. Cell viability was verified in parallel using CellTiter-Glo®.

## 4. Conclusion

Through SAR exploration, we successfully optimized the low-potency hit **E14** (KD = 138 µM) into the lead compound, **Z17** which represents a 20-fold improvement in affinity. SPR analysis confirmed that **Z17** binds potently to human CHI3L1 (KD = 6.0 µM). Additionally, specificity assay showed that **Z17** exhibits high selectivity toward CHI3L1 while showing no detectable binding to seven other neuroinflammatory proteins which is a marked improvement over the non-specific reference compound **G28**. **Z17** reinstated crucial astrocytic function by rescuing CHI3L1-induced deficits in A*β* uptake, restoring lysosomal proteolytic activity and normalizing lysosomal pH homeostasis in human iPSC-derived astrocytes. Furthermore, the *in vitro* pharmacokinetic (PK) evaluation demonstrated that **Z17** possesses a favorable profile for central nervous system (CNS) development which was characterized by balanced lipophilicity (LogD7.4 = 2.39 ± 0.22), high passive permeability (4.6 × 10⁻⁶ cm s⁻¹) and a wide safety margin against hERG inhibition and cytotoxicity. These combined findings establish **Z17** as a selective CHI3L1 inhibitor capable of simultaneously mitigating neuroinflammation and restoring astrocytic clearance mechanisms, positioning it as a highly promising therapeutic candidate for Alzheimer’s disease.

## Acknowledgments

This work was supported by the National Institute of Neurological Disorders and Stroke under grant number R01NS136524 (PI: Gabr).

## Data availability

Data will be made available on request.

