## Supporting Information for "Restoring Amyloid Clearance via Astrocytes: Z17 Is a Selective Inhibitor of CHI3L1 in Alzheimers Disease"

* To whom correspondence should be addressed:

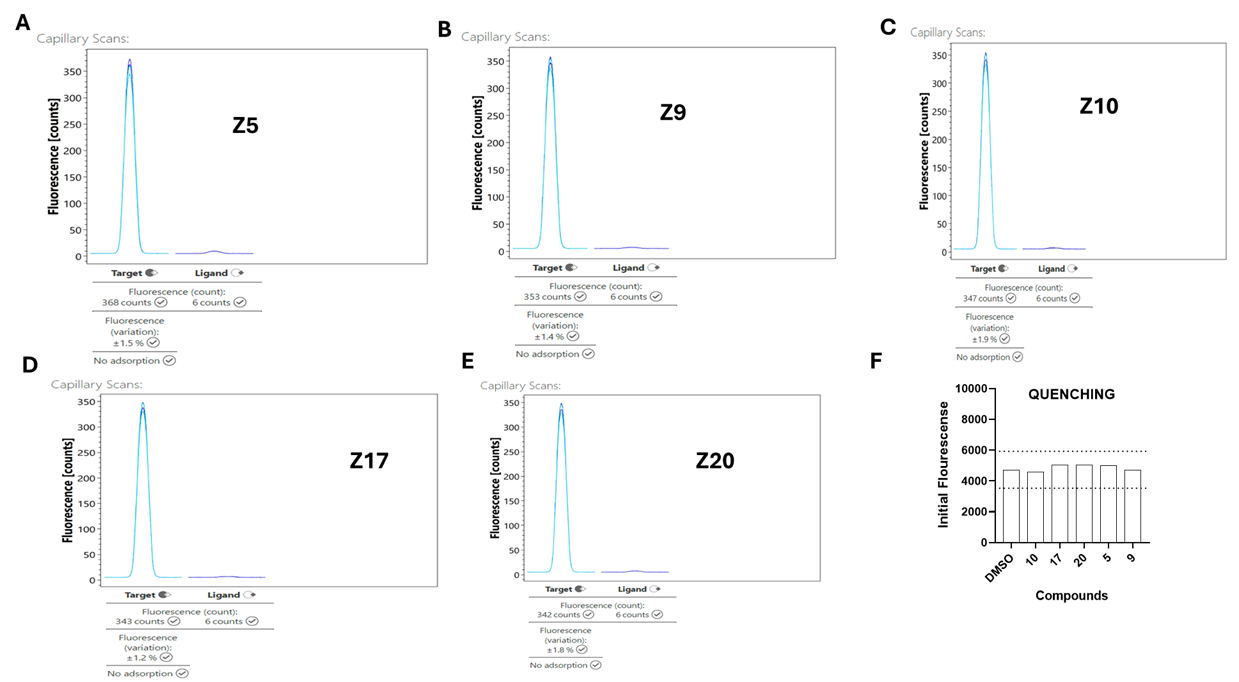

**Figure S1.** Assessment of potential artifacts from autofluorescence and quenching in MST assays. (A–E) Autofluorescence of test compounds was evaluated by comparing the fluorescence of labeled protein in 2.5% DMSO with buffer containing 250 μM compound in 2.5% DMSO. (F) Quenching was assessed by comparing fluorescence of 20 nM dye in buffer with DMSO to that of 20 nM dye incubated with each compound. None of the tested compounds showed significant autofluorescence or quenching under these conditions.
